# Measurement of ocular transmission in living human eyes with a double-pass system

**DOI:** 10.1101/606640

**Authors:** Roberto F. Sánchez, Aníbal G. de Paul, Francisco J. Burgos-Fernández, Meritxell Vilaseca, Jaume Pujol, Luis A. Issolio

## Abstract

**Purpose:** To develop a methodology based on a double-pass system to obtain useful information about the transmission of ocular media, performing noninvasive measures *in vivo*.

**Methods:** This noninvasive procedure consists of recording double-pass images at different voltages of a laser diode of 780 nm and the determination of the scattering in an area between 25 and 35 arc minutes of each image.

**Results:** Ocular scattering showed a linear behavior respect to the voltage of the laser and the slope of the linear fit was proportional to the transmittance squared of the media evaluated. The relationship between the ocular light scattering of the images and the transmittance values of several filters located into an artificial eye was used as a calibration function. The measurements performed in a group of ten subjects with ages between 25 and 45 years old presented a mean direct transmittance of the whole eye including retina of 42.7 %, which agrees with the bibliography. No differences between dark eyes and light eyes were found.

**Conclusion:** We have developed a method to determine the transmittance of the human eye in vivo for a wavelength of 780 nm using the double-pass method, commonly used for the determination of the optical quality of an eye.

## Introduction

Knowledge of transmission of the eye can be useful both for clinical applications where it is important to estimate the amount of light actually reaching the photoreceptors and in the field of illumination where it is often necessary to characterize visual stimuli not only with luminance, which represents the light arriving the eyes but also with retinal illuminance, which includes in its rigorous definition the transmittance of the eye [1].

Boettner and Wolter, as well as Geeraets and Berry, have measured the direct and total transmittance of isolated components of the ocular media, including the cornea, aqueous, lens and vitreous, but without including retina; using enucleated eyes of human donors and rhesus monkey [2-4]. They considered a wide range of wavelengths including visible, near ultraviolet (UV) and near infrared. Those results were confirmed by Alpern et al. with *in vivo* measurements performed on subjects with structural abnormalities in the fundus of the eye (chorioretinal coloboma). That condition allowed the determination of the amount of light transmitted by the eye, through a direct comparison of the component reflected in the sclera of a light beam with a reference [5]. Dillon et al. proposed an invasive method for measuring the absorption spectrum of the anterior segment of the intact eye. The true spectrum of light that is transmitted to the retina was calculated for four different species that are commonly used in animal model experiments [6]. Van Norren and Vos have estimated the spectral transmittance of the eye as the measured difference in the scotopic visual sensitivity of two wavelengths with the same rhodopsin absorbance [7]. Many other studies of the human ocular media were limited to a single component of the eye, usually cornea o lens. In 2007, van de Kraats and van Norren proposed empirical equations to estimate changes in ocular transmittance as a function of eye age [8]. Based on existing data and these equations, the International Commission on Illumination (CIE) established the transmission of the standard observer for UV, visible and infrared which can be used as a reference [9]. Currently, there is no simple procedure for measuring the transmittance of the whole eye of a healthy subject.

The purpose of this work is to take advantage of the double-pass technique to estimate in a direct way the transmittance of the eye at a given wavelength. The double-pass method is a simple, fast, safe, and non-invasive technique to obtain information about the energy that enters the eye, crosses the ocular media, reflects in the fundus and returns. This technique is based on imaging a point source on the retina, and then recording the reflected light through a CCD camera [10]. The acquired image is the autocorrelation of the point spread function (PSF) of the eye [11], which describes how the optical system behaves against a point light source [12].

It is known that in the retina, light is reflected by different layers and a significant portion of the light reflected by the background comes from the choroid and is dependent on pigmentation and wavelength [13,14]. In a doubled-pass system, the imaging formation approximates that of a confocal system, so that most of the light coming from the deeper layers not conjugated to the camera are spreading, adding a background signal to the recorded image. Thus, the double-pass image shows a very narrow peak mounted on a fairly wide tail [15]. Scattering affects both the peak and the wide-angle part of the PSF, unlike aberrations, which are limited to the very narrow-angle part of the PSF [16-18]. Several authors showed that it is possible to obtain information about intraocular scattering by analyzing the peripheral zone of the double-pass image up to 20 arcminutes [19-21].

Our goal is to develop a methodology based on a double-pass system to measure the transmission of ocular media, performing noninvasive measures *in vivo*. In this work, we propose a simple method to determine the direct transmittance of the eye including the retina, based on the acquisition and analysis of double-pass images.

## Material and Methods

### Experimental setup

A scheme of the optical system mounted is shown in Fig 1. A point source (O) from a 780 nm laser diode (MC7850CPWR-SMF) coupled by single-mode optical fiber to a collimator lens L1 (f = 15 mm) is projected onto the retina (O’). After reflecting on the retina and a double pass through the ocular media, a CCD camera records the double-pass image (O”). The diaphragm PE (Ø = 2 mm) is conjugated to the pupil plane of the eye and acts as the effective input pupil of the system. After being reflected by beam splitter BS1 and mirror M1, the beam passes through the Badal system formed by the lenses L2-L3 (f = 200 mm) and by the mirrors M2-M3 that regulate beam vergence, allowing correction of refractive errors such as myopia or hyperopia of the eye before performing the measurement. After reflection in mirror M4, the eye forms the image of the point source on the retina. A second diaphragm PS (Ø = 4 mm), located after the beam splitter (BS2), is also conjugated to the pupil of the eye and acts as the effective exit pupil (provided the natural pupil of the eye is greater than PS). Then a lens L4 (f = 100 mm) forms the double-pass retinal image in a CCD camera (UI-2220ME-M, 8 bits, 768×576 pixels) integrating light from the retina during the set exposure time. The subject’s head is placed in a chin rest which allows the correct centering and control of natural pupil with respect to the artificial one. A camera CMOS (UI-1221LE-M-GL) and the lens L5 (f = 50 mm) are mounted to ease this process. The fixation of the eye is facilitated using a fixation target (FT) located at the optical infinity by the lens L6 (f = 35 mm).

**Fig 1.**
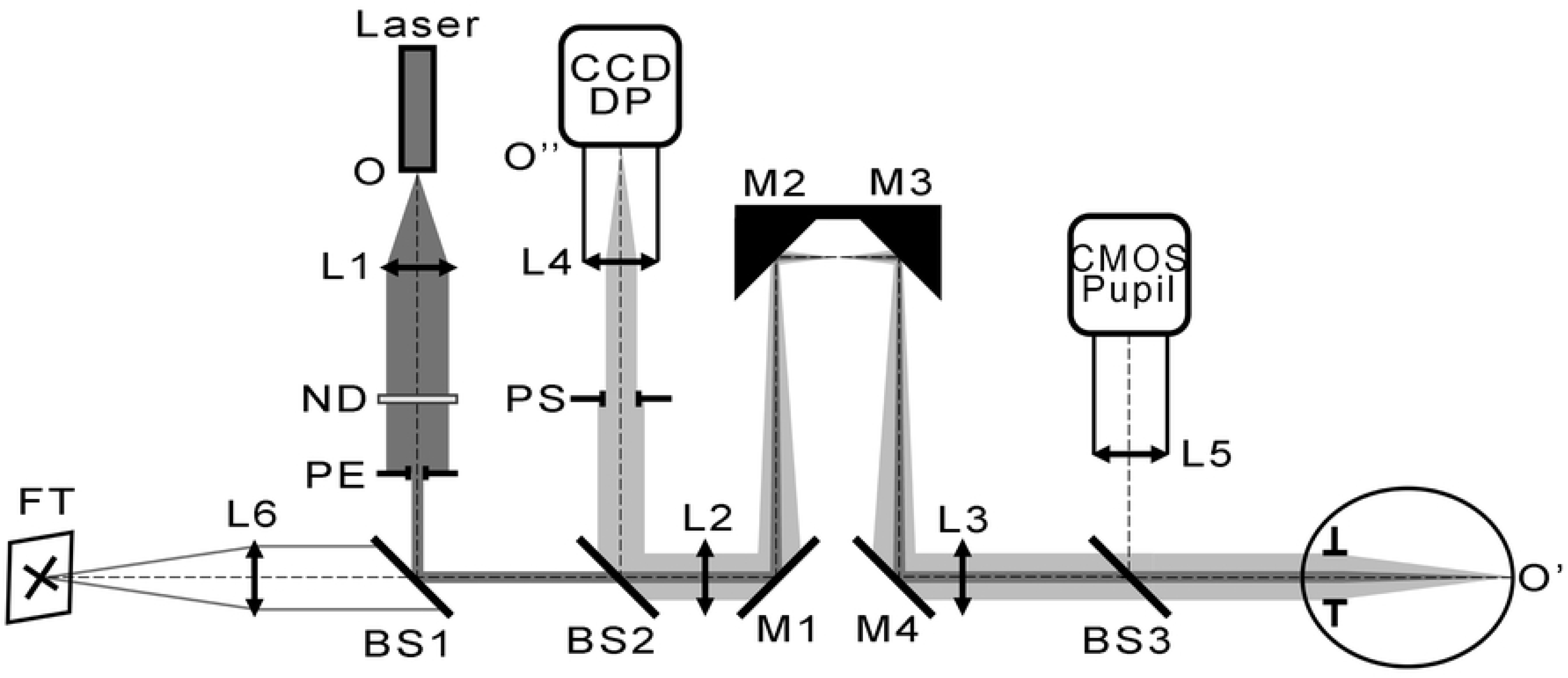
Schematic representation of the double-pass setup mounted for this work. See text for further details.

To eliminate back reflections from lenses L2 and L3, these elements were tilted slightly, so that the specular reflection at the interfaces of these lenses was deflected off the field picked up by the CCD sensor. The corneal reflex and the lens reflex (Purkinje images) are not observed in the double-pass image captured, due to a small decentering of the measuring beam with respect to the pupil of the subject. Other diffuse reflections are reduced by light traps (not shown in Fig 1). Finally, there are still spurious reflexes that are integrated by the sensor during the recording of the images. However, these are considered during postprocessing by subtracting the captured background when removing the eye.

### Methodological proposal

The proposed procedure to determine the transmission of the ocular media consists of recording a series of double-pass images, each taken at a different intensity of the laser diode, which was controlled by a supply voltage from 0 mV to 4000 mV. The intensity in each record was increased from low values, where the maximum of the image did not saturate, until values high enough to obtain images with the greater gray level possible in the more peripheral pixels. In the capture performed at each intensity an image of the background was also obtained, which was then subtracted from the corresponding double-pass image. The centroid of each image was calculated and, taking this position as a center, the radial profile was obtained (averaged of pixels at different radii); from which the gray levels of an area between 25 and 35 arc minutes were averaged. This average, which we will call hereafter double-pass scattering (DPS), was determined for each image taken with different levels of the voltage of the laser (LV).

If the low order aberrations are corrected during the acquisition, then the central part of the double-pass PSF is affected by higher order aberrations and light scatter. However, the area of the PSF analyzed in our work is only affected by light diffused in the fundus and light scattered in the ocular media.

In Fig 2, DPS values are plotted as a function of LV and a linear relationship between them is observed. A straight line can be fitted to the data using least squares.

**Fig 2.**
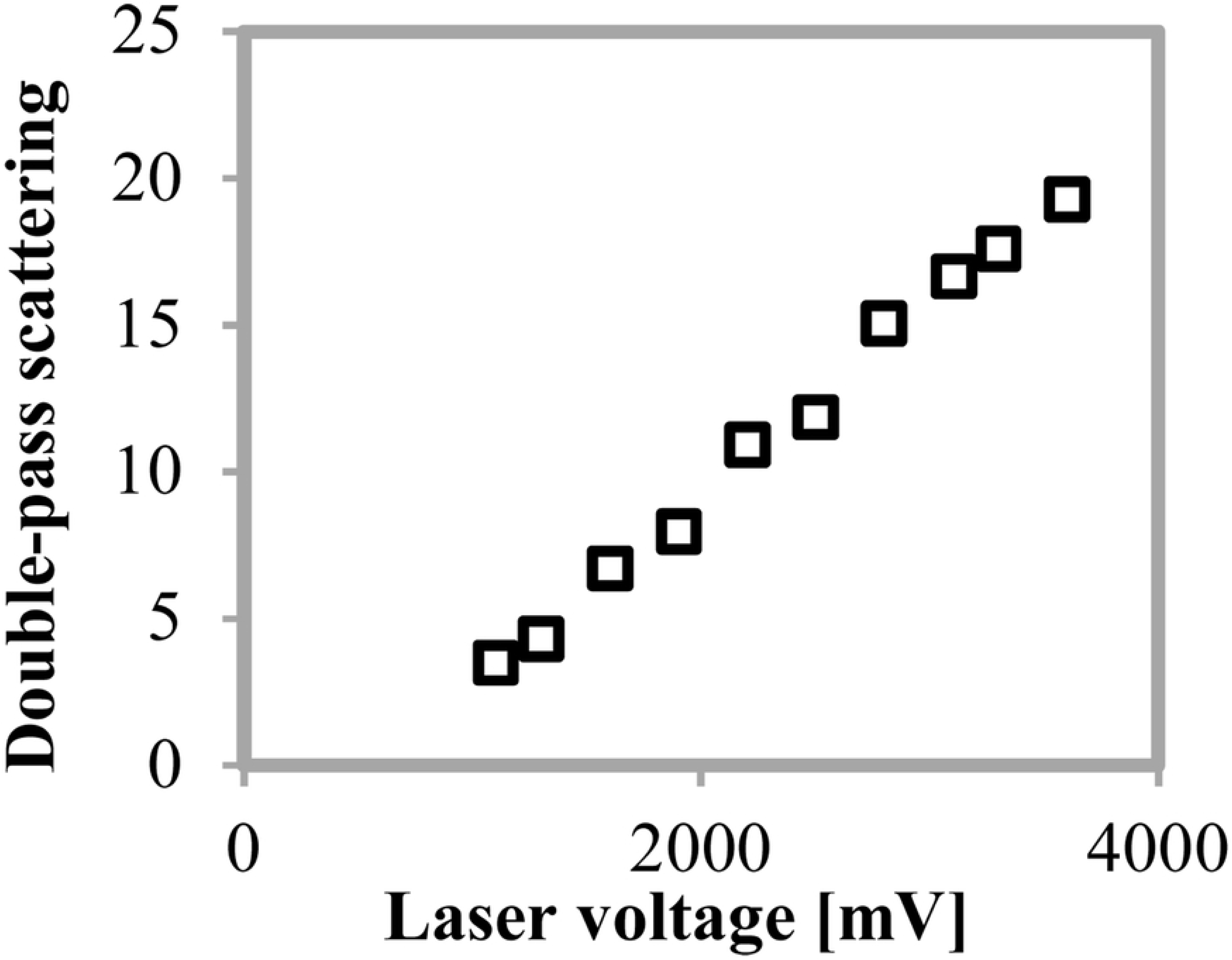
A typical curve of double-pass scattering (DPS) as a function of the laser voltage. DPS was computed as a mean of the grey levels in a ring of 25 to 35 arc minute of the double-pass image.

Our hypothesis was that the scattering increases proportionally with the intensity of the laser diode, not affecting the slope. Moreover, as the light has to cross the ocular media twice, we expect the slope to be proportional to the transmission squared of the eye.

### Measurements in an artificial eye with filters

To test the proposed methodology, measurements were made with an artificial eye built in the laboratory consisting of two lenses simulating the cornea (f = 25 mm) and the lens (f = 50 mm), plus a black diffuser screen located in the focal plane of the lens system acting as a reflective retina. In order to simulate eyes with several transmissions, neutral filters of different optical density (0.1, 0.3, and 0.5) were placed in front of the artificial eye. Additionally, measurements with photography effect filters (BPM1, BPM2, and C3020) that have been shown to be very suitable to simulate the same type of scattering produced by a cataract eye were performed [22,23].

Previously, these filters were characterized by their direct transmittance and those with transmission values that allowed to achieve laser energy levels comparable to those used in double-pass measurements in real eyes were selected. The transmittance measurement was performed using the 780 nm laser as an illuminator and a sensor (E2V, Spindler & Hoyer) to record the amount of luminous flux emitted by the source as well as the amount of light passing through each filter. We placed the detector further from the filter and used an aperture before the detector which subtended an angle at the filter of 1°. Thus, the sensor detected only that part of the transmitted radiation which exited the filter within a one-degree angle collinear with the incident radiation. From this data, we computed the direct transmittance of the filter. This component is also called the focusable transmittance and is important for understanding the deposition of laser energy on the retina.

Measured transmittances at 780 nm in each of the filters used to obtain the calibration curve with the artificial eye are shown in Table 1.

**Table 1.** Direct Transmission Measurements.

| Filter | Measured Transmission |
| --- | --- |
| ND 0.01 | 97 % |
| ND 0.1 | 88 % |
| ND 0.3 | 56 % |
| ND 0.5 | 34 % |
| BPM1 | 68 % |
| BPM2 | 59 % |
| C3020 | 85 % |

In Fig 3 are presented the profiles of a set of double-pass images taken in the artificial eye without any filter. As the voltage of the laser increases, the central area of the curve saturates and a measurable variation in the peripheral zone (the skirt of the curve) can be observed.

**Fig 3.**
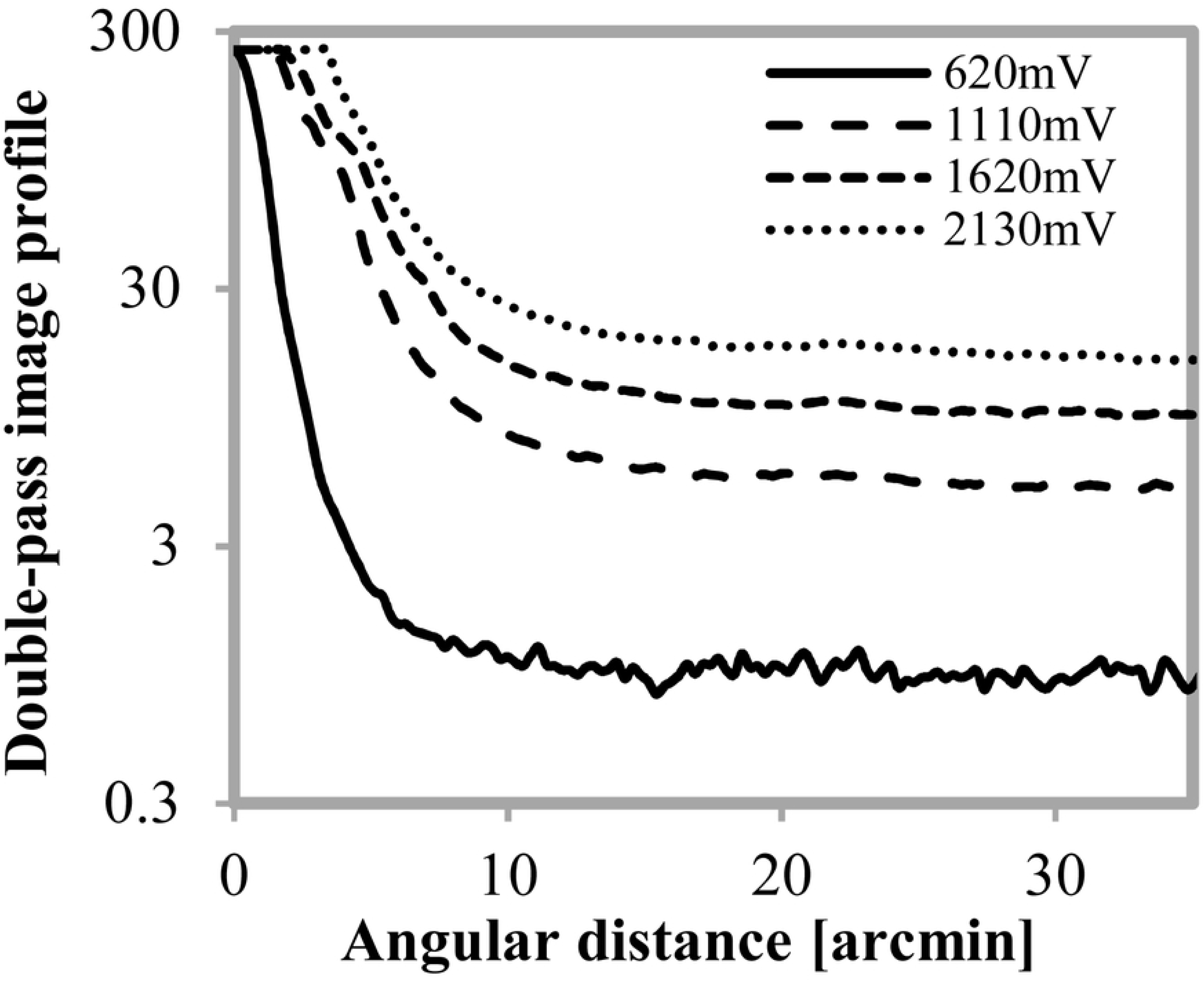
Double-pass image profiles obtained in the artificial eye without any filter. The curves correspond to four different voltage applied to the laser diode (620, 1110, 1620, and 2130 mV).

In Fig 4 are plotted the DPS values as a function of laser voltage for each of the considered filters, as well as for the artificial eye without a filter. Straight lines were fitted for each condition and the slopes were obtained. As the transmission of the filters increases, the slopes are ordered from lowest to highest, which leads us to call them ocular transmission index (OTI). Plotting the relationship between OTI and the direct transmittance measured at each filter, a quadratic curve (Fig 5) was obtained and then used to determine the transmission of the ocular media in subjects.

**Fig 4.**
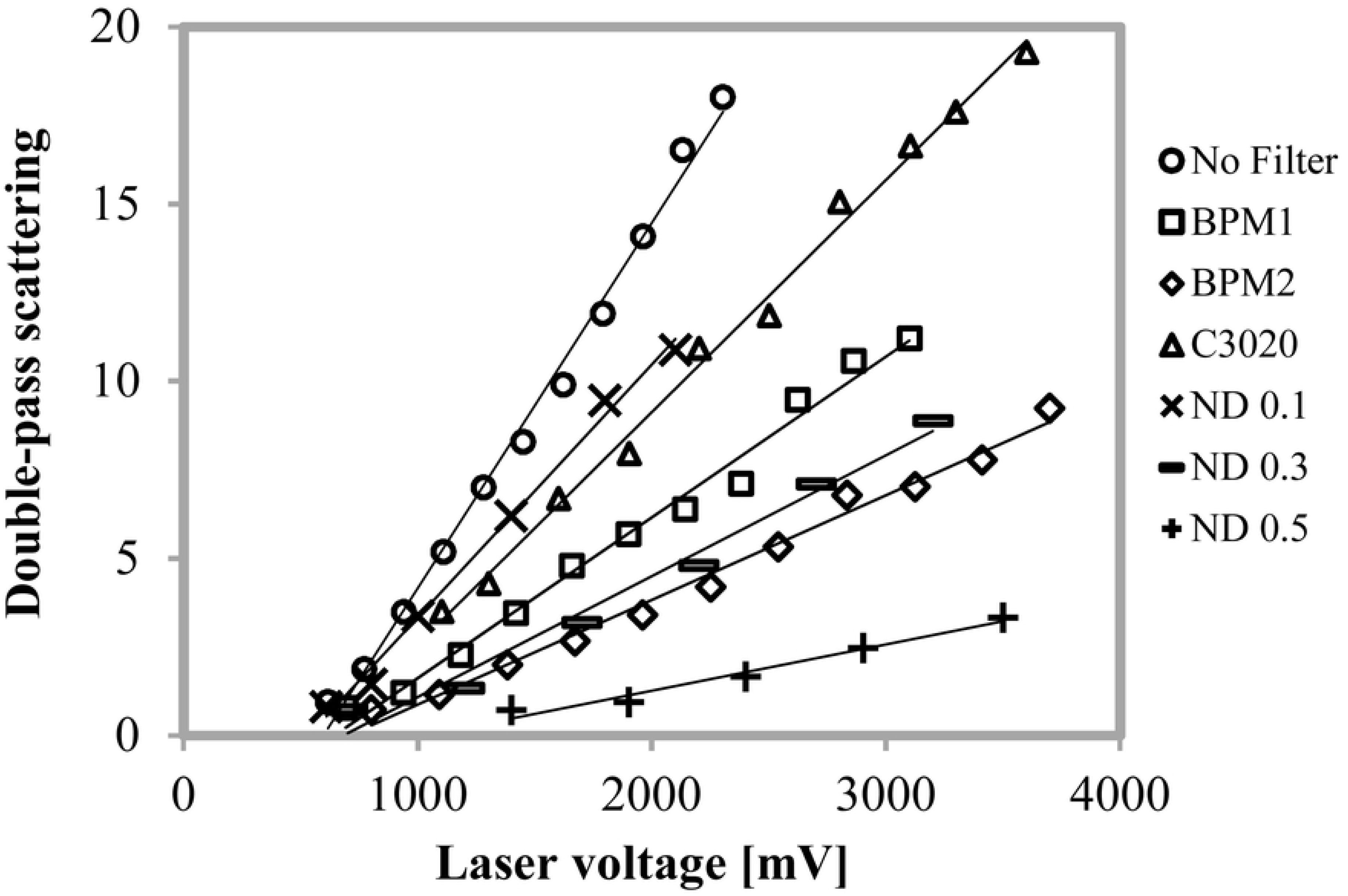
DPS as a function of laser voltage obtained in the artificial eye with and without filters. Three photography filters (BPM1, BPM2, and C3020) and three neutral density filters (ND 0.1, ND 0.2, and ND 0.3) were used along with the artificial eye. The straight lines correspond to the linear fit performed.

**Fig 5.**
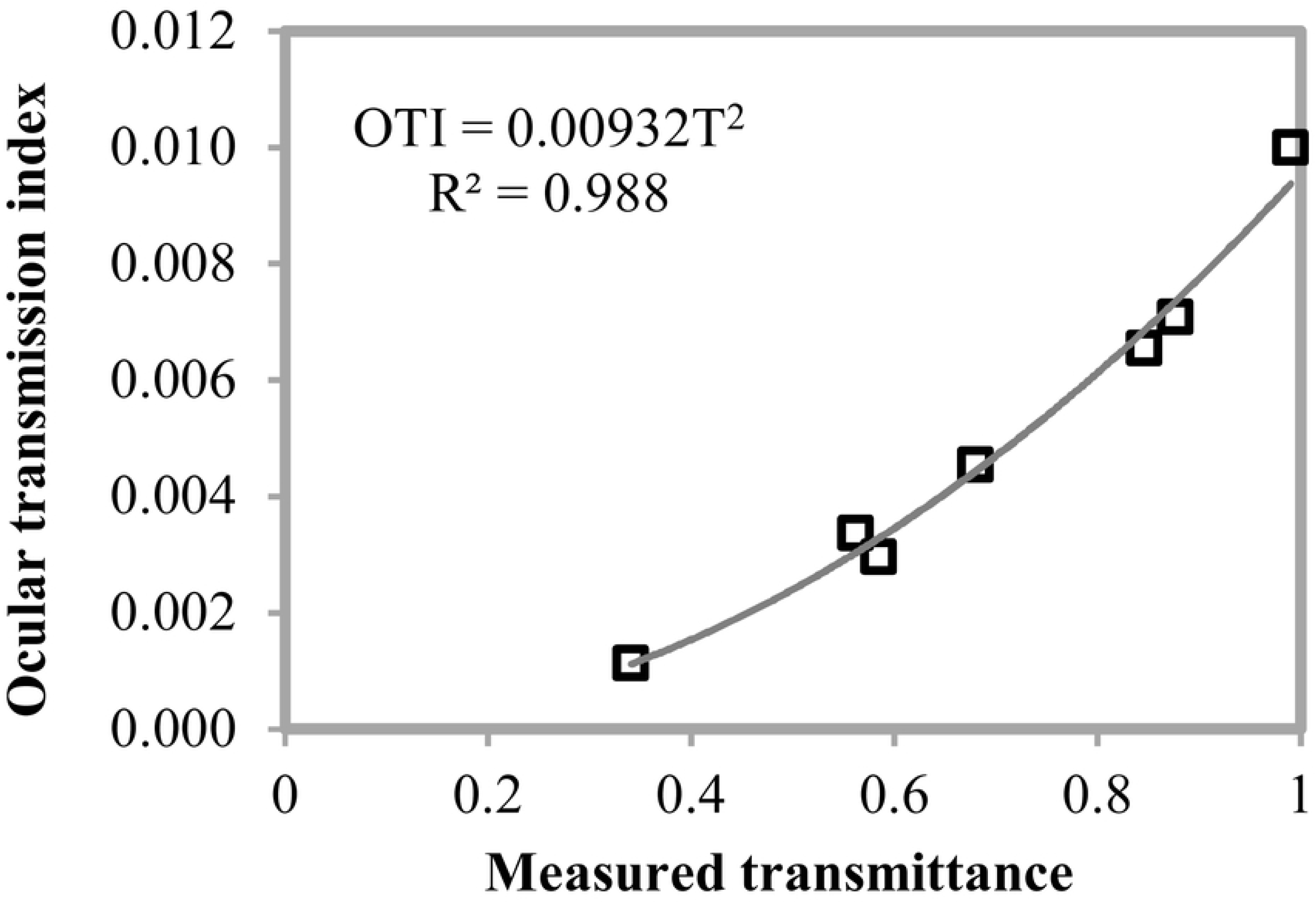
Ocular transmission index (OTI) as a function of the transmittance measured in the filters. OTI values are the slopes of the straight lines in Fig 4.

From the curve fitted to the data plotted in Fig 5 can be deduced the following expression to estimate the direct transmission of ocular media (*T*) as a function of the OTI:

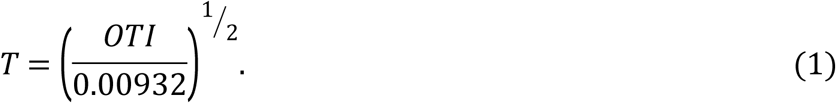

### Reflectance analysis

The energy captured in the double-pass image could also depend on the fundus reflectance of the human eye or the reflective surface of the artificial eye. To analyze the effect of this factor on the measurement, double-pass images of the artificial eye with different reflective surfaces (white paper and black paper) were taken. The results presented in Fig 6 (a) show that although very different profiles of double-pass images are obtained in each condition, there are no significant changes in the peripheral zone of the curves (note that the vertical axis is in logarithmic scale), which can also be seen in Fig 6 (b) where the fitted curves do not change when the reflectance in the artificial eye is modified (independent sample t-test was used for comparing means in each condition, P-value > 0.05 in all cases).

**Fig 6.**
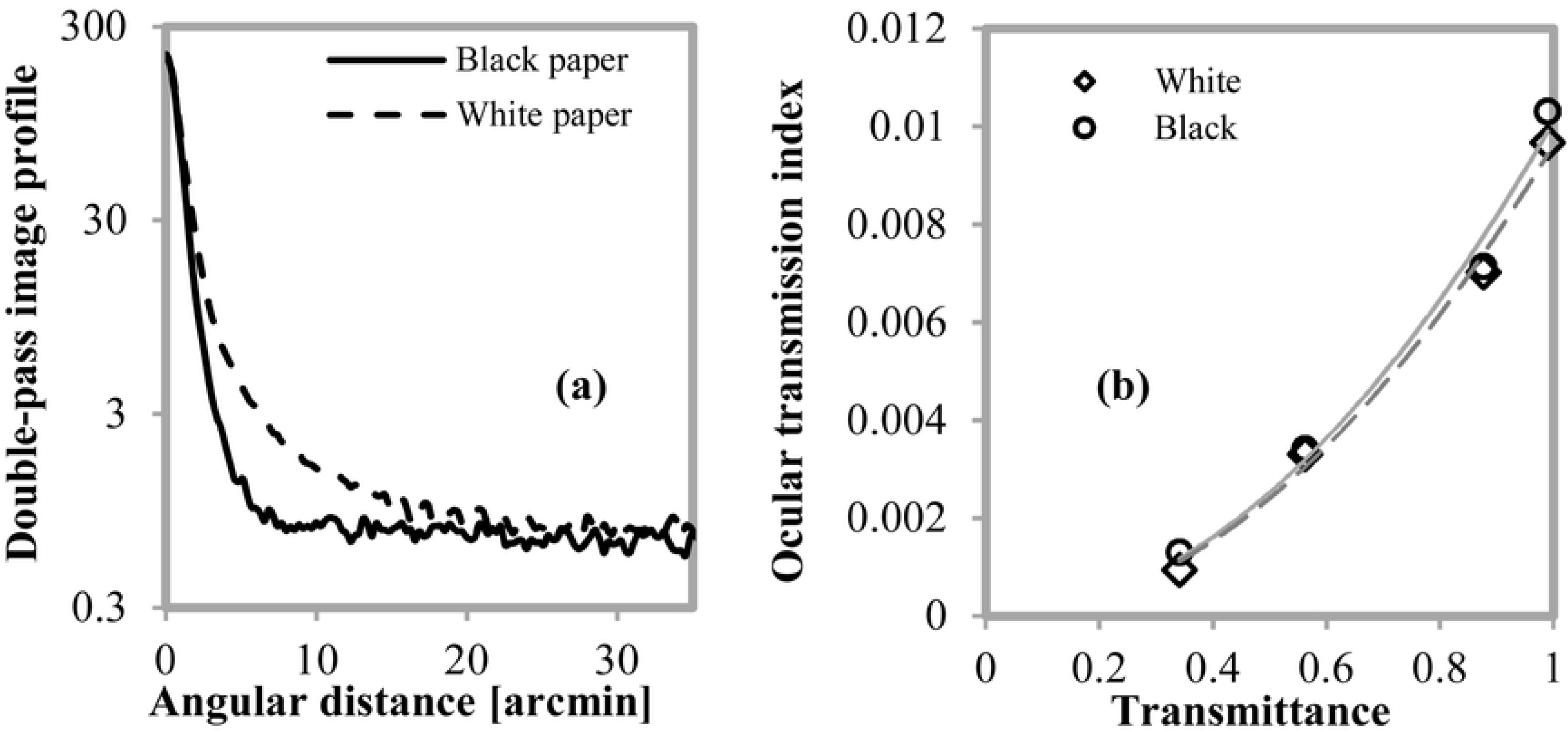
Analysis of the effect of reflectance on the measurement of OTI. (a) Double-pass image profile obtained for two different papers used as artificial retinas (black and white) measured at the same laser voltage. (b) Curves fitted to the OTI data vs. transmittance for two papers used. Conditions measured are the three neutral filters (ND 0.1, ND 0.3, and ND 0.5) and the artificial eye without a filter. The solid line corresponds to black paper and dashed line to white paper.

In order to analyze the OTI independence regarding the variation of the reflectance between different retinas, five subjects were evaluated. The set of curves were obtained considering the same intensity of the laser provided by a voltage of 1400 mV.

Fig 7 shows the double-pass image profiles determined in this group of subjects. Values in the zone between 25 and 35 minutes of arc vary around a gray level of 1.0 ± 0.3 for all the subjects, which verifies the hypothesis about the independence of the OTI respect to the reflectance of the retina.

**Fig 7.**
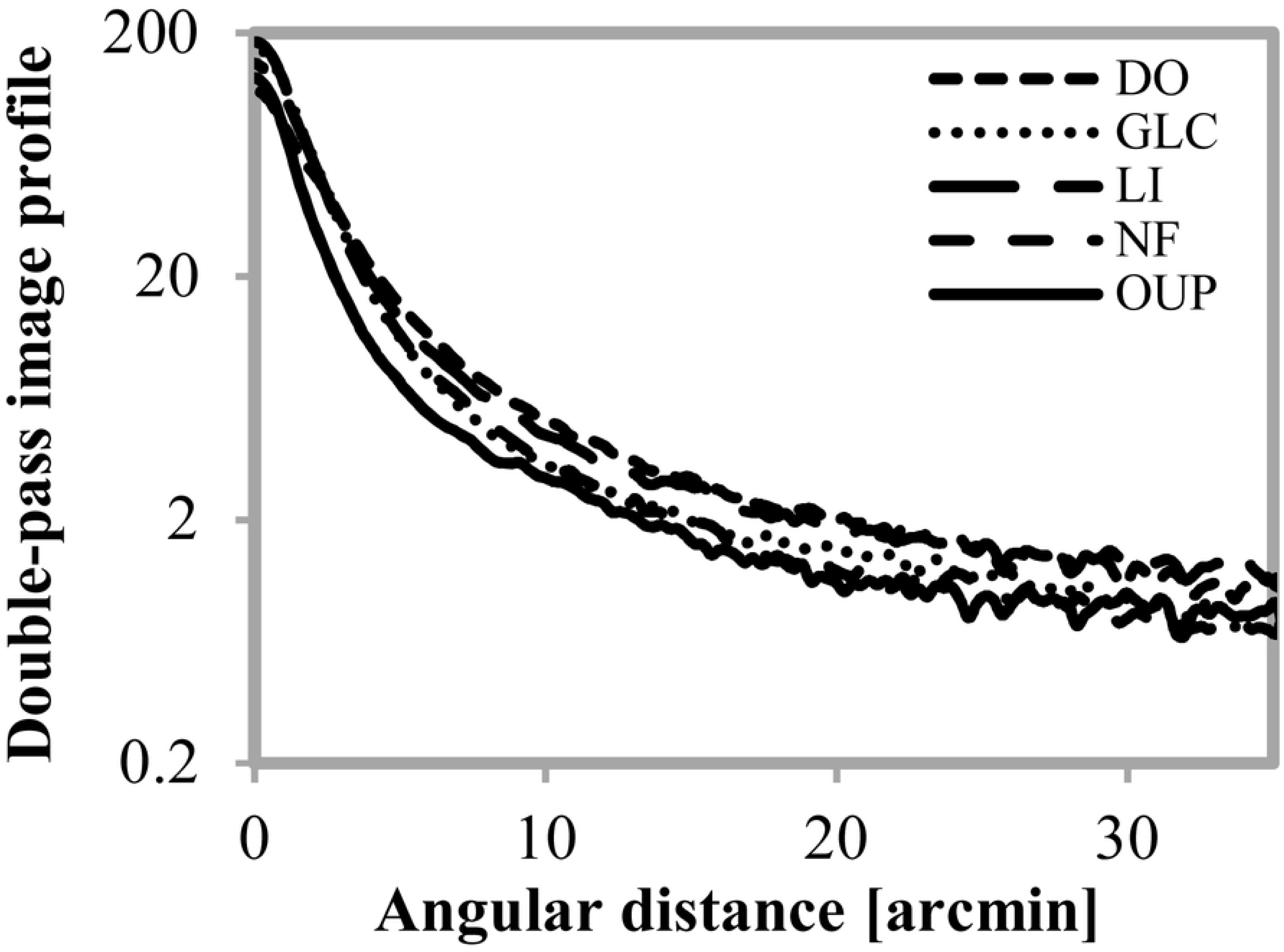
Double-pass images profiles determined in five subjects with the same laser voltage.

### Measurements in volunteers

Once the method was tested on the artificial eye with different filters and the calibration curve was determined (Fig 5), double-pass images were recorded in eyes of ten volunteers aged 25-45 years (Group A). Seven of the subjects were classified as dark-eyed (four dark brown and three light brown) and three as light-eyed (blue), according to a classification by simple iris observation [24]. Depending on the characteristics of the eye, intensities between 1000 and 4000 mV were used and at least five DPS were measured in each subject to ensure a good fit. In each case, the dominant eye was chosen to perform the measurement.

Additional measurements were made in another group of five volunteers (Group B) to evaluate the range of low transmittance values. For this set of measurements, neutral density filters (0.01, 0.1, and 0.3) were placed in front of the eye of each subject and then the OTI slope was obtained for each condition. For all measurements, the exposure levels were never greater than the maximum permissible exposure (14.45 W/m^2^) which was established by the current standard regulating the use of laser radiation in living tissue [25].

### Ethical considerations

Ethical approval for this study was obtained from the Comité de Ética en Investigación (CEI) of the Universidad Nacional de Tucumán and Consejo Nacional de Investigaciones Científicas y Técnicas (RESOLUCION N° 26/2018). All the subjects were informed about the object of the study, and a written informed consent was obtained, following the tenets of the Declaration of Helsinki.

## Results

In Fig 8 are presented DPS data as a function of laser voltage for Group A, and the straight lines fitted in all the subjects. All the data show a good linear fit, with an R^2^ always greater than 0.85.

**Fig 8.**
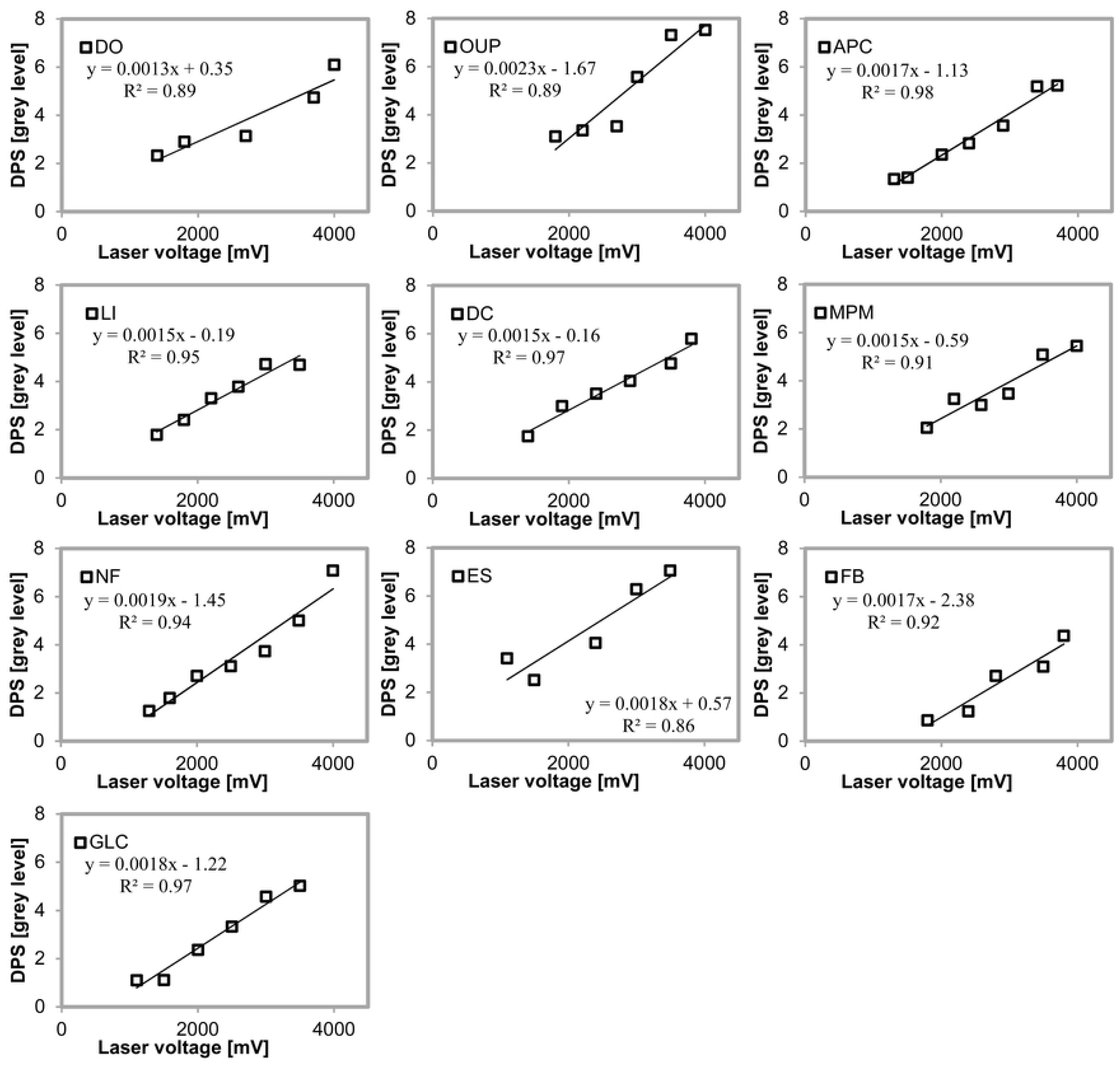
Double-pass scattering expressed in terms of grey level as a function of the power supply voltage of the laser diode. For each subject, the fitted straight line and the coefficient of determination are shown.

Table 2 presents the OTI values determined in each subject and the direct transmittance value computed from Eq 1.

**Table 2.** Ocular transmission index and direct transmission of the eye for each subject of Group A.

| Subject (iris color) | OTI | Direct transmittance of the eye % |
| --- | --- | --- |
| NF (dark) | 0.00194 | 45.6 |
| MPM (dark) | 0.00151 | 40.3 |
| OUP (dark) | 0.00235 | 50.2 |
| GLC (dark) | 0.00183 | 44.3 |
| DO (dark) | 0.00128 | 37.1 |
| FB (dark) | 0.00169 | 42.6 |
| APC (dark) | 0.00173 | 43.1 |
| LI (light) | 0.00150 | 40.1 |
| ES (light) | 0.00178 | 43.7 |
| DC (light) | 0.00149 | 40.0 |

Pigmentation (iris color) is shown in parentheses. OTI, ocular transmission index.

The obtained values ranged from 37 % to 50 %, with a mean value of 42.7 ± 3.6 % (Mean ± SD) for the sample considered. The mean value for subjects with light eyes was 41.3 ± 2.1 % (Mean ± SD), whereas for subjects with dark eyes the mean was 43.3 ± 4.1 % (Mean ± SD). There is no significant difference in the transmittance found for these groups (P-value = 0.335).

Fig 9 (a) shows the DPS values for the four conditions evaluated in one of the subjects of the Group B (CT without a filter, CT + ND 0.01, CT + ND 0.1, and CT + ND 0.3) and the performed fit. From the slopes of these lines, the transmittance of each condition (eye or eye plus ND filter) was estimated using Eq 1 (measured transmission). The transmittance of each condition can be predicted (computed transmission) by multiplying the known transmittance of each filter (Table 1) by the transmittance measured in the eye without filters. All the subjects from Group B showed a similar pattern of results.

**Fig 9.**
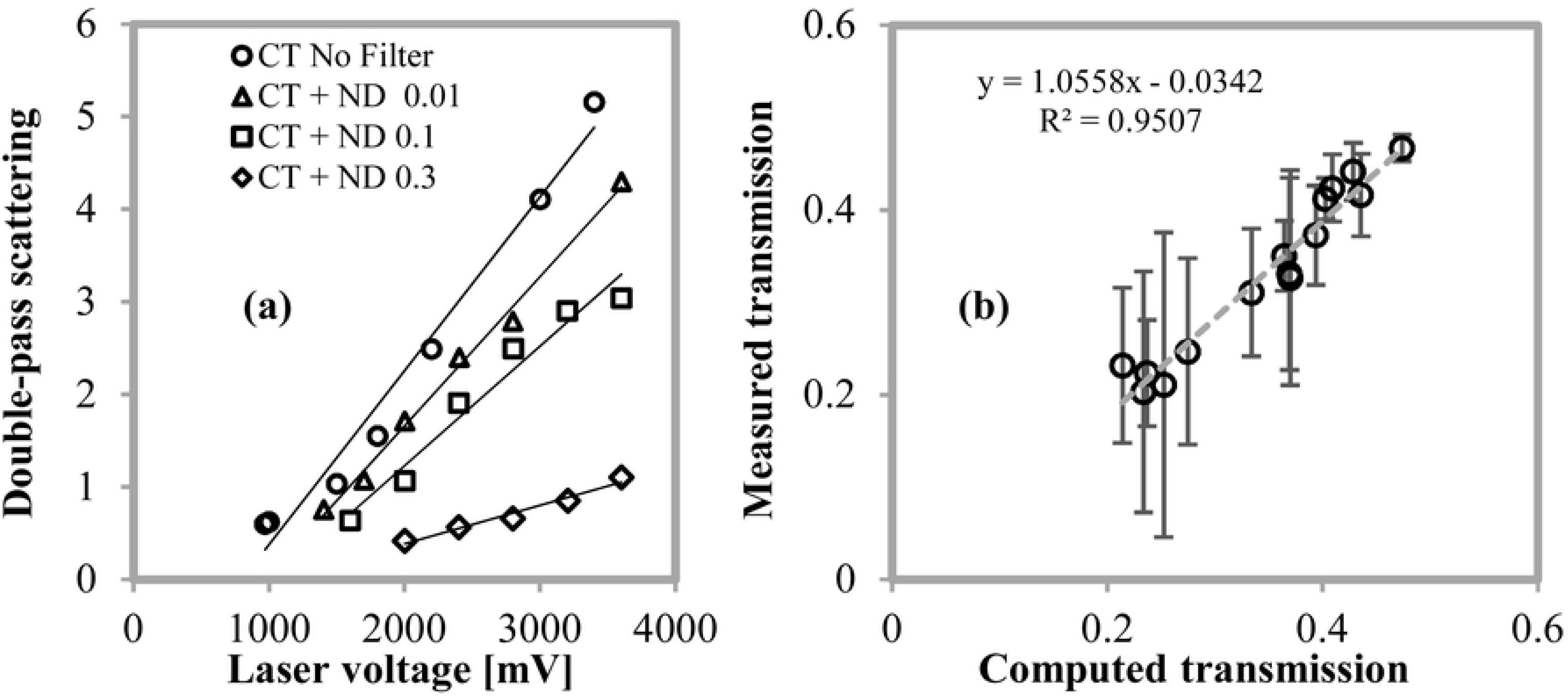
Measured transmission vs predicted transmission. (a) DPS as a function of laser voltage obtained in the subject CT with different neutral filters and the condition without a filter. (b) Direct transmission measured in each filter along with the eye for the five volunteers vs the computed transmission. Error bars show the relative error with respect to the expected value.

Fig 9 (b) shows the transmittance values measured in the five subjects with three filters, as a function of the computed transmittance. The high correlation showed in Fig 9 (b) was expected and adds consistency to our results. However, it can be seen that errors grow for low transmittances (relative errors become 15% for transmittances of 0.2).

## Discussion

This is the first article proposing a methodology to measure the *in vivo* direct transmittance of the whole eye, that is, including the retina. Boettner reported 54 % for direct transmittance at 780 nm in eyes corresponding to a similar range of age to that used in our work [3]. This value corresponds to the transmission of pre-retinal media (cornea, aqueous humor, lens, and vitreous). Moreover, Boettner’s work shows a measure of the transmittance of rhesus monkey retina, which is considered anatomically similar to that of the human eye, of 80 % for a wavelength of 800 nm, which means a total transmittance of 43 %. Our results, based on double-pass measurements that take the image reflected in the inner layers of the retina, provide an estimate of the direct transmittance of the eye including retina of 42.7 %. There is no significant difference between our result and those reported in previous works (P-value = 0.675).

As with other optical quality measurements derived from the double-pass method [26], our measurement of transmittance is affected by uncorrected low-order aberrations (defocus and astigmatism). In our measurements, the refractive errors of the subjects were corrected with the spherical equivalent by means of the Badal system, therefore, in some subjects, there could be some uncorrected astigmatism effect that adds error to the measure.

Between each capture, from the beginning of the experiment until the end a few minutes pass, therefore, eye movements were inevitable. This would cause the laser to strike a different area of the retina with a slightly different reflectance, however, those variations did not produce changes on the measured slope. In that sense, the behavior of the eye was similar to that presented by the artificial eye (Fig 6).

Ginis et al. [24] have reported differences in a small sample in the straylight measured in subjects with light eyes and dark eyes. Ginis’s measurements show that diffuse light from the fundus contributes significantly to the total straylight for wavelengths longer than 600 nm, especially in the measurements made for small-angles of the PSF (30 arc minutes). In our study, no significant differences were found between the transmittances measured in subjects with different pigmentation.

According to Fig 9, the technique described in this work can be used for a wide range, however, because low values of transmission have a significant error the method would not be suitable for this range. For this, the present characteristic of the sensor limits the scope of the proposed method due to the low energy levels that reach it in the evaluated region. However, since we are only interested in the energy reaching this area and not in the whole PSF, single-pixel sensors could be used instead of a CCD to increase the dynamic range and capture much lower levels.

The transmittance values found to correspond to the wavelength of 780 nm and it is possible to determine the transmittance for other wavelengths using the same method, considering the modification in the sensor previously mentioned. In addition, a double-pass system with a polychromatic source has been developed in recent years [27] and the spectral transmittance of the human eye could be also determined based on the same configuration and the procedure described here. The spectral transmittance can be useful for specific applications such as in the study of intrinsically photosensitive retinal ganglion cells using a five-primary photostimulator, where it is necessary to characterize the individual differences in pre-receptoral filtering in each subject [28,29].

## Conclusion

We have developed a procedure to determine the transmittance of the human eye *in vivo* for a wavelength of 780 nm using the double-pass method, commonly used for the determination of the optical quality of an eye. The process requires taking double-pass images with different laser beam intensities and determining the slope of the straight line fitted to the obtained data. From this value and using a calibration equation we describe here; the direct transmittance of the eye is obtained.

## Acknowledgments

We thank Jose Barraza, Ph.D., UNT-CONICET for the assistance and for comments that greatly improved the manuscript.

## References

1. Le Grand Y. Light, Colour and Vision. Second English edition, translated from the French by Hunt RWG, Walsh JWT, and Hunt FRW. Chapman and Hall, London, 1968.

2. Boettner EA, Wolter JR. Transmission of the ocular media. Invest Ophthalmol Vis Sci. 1962 Dec;1(6):776–83.

3. Boettner EA. Spectral transmission of the eye. Final contract report AF41(609)-2966, University of Michigan, Ann Arbor MI, July 1967. [Available from NTIS as AD 663246].

4. Geeraets WJ, Berry ER. Ocular spectral characteristics as related to hazards from lasers and other light sources. Am J Ophthalmol. 1968 Jul;66(1):15–20.

5. Alpern M, Thompson S, Lee MS. Spectral transmittance of visible light by the living human eye. J Opt Soc Am. 1965 Jun;55(6):723–7.

6. Dillon J, Zheng L, Merriam JC, Gaillard ER. Transmission spectra of light to the mammalian retina. Photochem Photobiol. 2000 Feb;71(2):225–9.

7. Norren DV, Vos JJ. Spectral transmission of the human ocular media. Vision Res. 1974 Nov;14(11):1237–44.

8. van de Kraats J, van Norren D. Optical density of the aging human ocular media in the visible and the UV. J Opt Soc Am A Opt Image Sci Vis. 2007 Jul;24(7):1842–57.

9. Lund DJ, Marshall J, Mellerio J, Okuno T, Schulmeister K, Sliney D, et al. A computerized approach to transmission and absorption characteristics of the human eye. In: CIE 203:2012. 2012, International Commission on Illumination.

10. Santamaría J, Artal P, Bescós J. Determination of the point spread function of human eyes using a hybrid optical-digital method. J Opt Soc Am A. 1987 Jun;4(6):1109–14.

11. Artal P, Marcos S, Navarro R, Williams DR. Odd aberrations and double-pass measurements of retinal image quality. J Opt Soc Am A Opt Image Sci Vis. 1995 Feb;12(2):195–201.

12. Guirao A, Lopez-Gil N, Artal P. Double-pass measurements of retinal image quality: a review of the theory, limitations, and results. In: Proceedings Vision Science and its Applications; 2000 Feb 11; Santa Fe, New Mexico United States. OSA Technical Digest (Optical Society of America, 2000), paper NW4.

13. López-Gil N, Artal P. Comparison of double-pass estimates of the retinal image quality obtained with green and near infrared light. J Opt Soc Am A Opt Image Sci Vis. 1997 May;14(5):961–71.

14. Ginis HS, Perez GM, Bueno JM, Pennos A, Artal P. Wavelength dependence of the ocular straylight. Invest Ophthalmol Vis Sci. 2013 May;54(5):3702–8.

15. van den Berg TJ, Franssen L, Coppens JE. Straylight in the human eye: testing objectivity and optical character of the psychophysical measurement. Ophthalmic Physiol Opt. 2009 May;29(3):345–50.

16. van den Berg TJ. Analysis of intraocular straylight, especially in relation to age. Optom Vis Sci. 1995 Feb;72(2):52–9.

17. Franssen L, Tabernero J, Coppens JE, van den Berg TJ. Pupil size and retinal straylight in the normal eye. Invest Ophthalmol Vis Sci. 2007 May;48(5):2375–82.

18. van den Berg TJ, Franssen L, Kruijt B, Coppens JE. History of ocular straylight measurement: A review. Z Med Phys. 2013 Feb;23(1):6–20.

19. Artal P, Benito A, Pérez GM, Alcón E, de Casas A, Pujol J, et al. An objective scatter index based on double-pass retinal images of a point source to classify cataracts. PLoS One. 2011 Feb 4;6(2):e16823.

20. Vilaseca M, Romero MJ, Arjona M, Luque SO, Ondategui JC, Salvador A, Güell JL, et al. Grading nuclear, cortical and posterior subcapsular cataracts using an objective scatter index measured with a double-pass system. Br J Ophthalmol. 2012 Sep;96(9):1204–10.

21. Paz Filgueira C, Sánchez RF, Issolio LA, Colombo EM. Straylight and visual quality on early nuclear and posterior subcapsular cataracts. Eye Res. 2016 Sep;41(9):1209–15.

22. Barrionuevo PA, Colombo EM, Vilaseca M, Pujol J, Issolio LA. Comparison between an objective and a psychophysical method for the evaluation of intraocular light scattering. J Opt Soc Am A Opt Image Sci Vis. 2012 Jul;29(7):1293–9.

23. de Wit GC, Franssen L, Coppens JE, van den Berg TJ. Simulating the straylight effects of cataracts. J Cataract Refract Surg. 2006 Feb;32(2):294–300.

24. Christaras D, Ginis H, Artal P. Spatial properties of fundus reflectance and red-green relative spectral sensitivity. J Opt Soc Am A Opt Image Sci Vis. 2015 Sep;32(9):1723–8.

25. Safety of Laser Products, IEC 60825-1 Standard (2007).

26. Martínez-Roda JA, Vilaseca M, Ondategui JC, Giner A, Burgos FJ, Cardona G, et al. Optical quality and intraocular scattering in a healthy young population. Clin Exp Optom. 2011 Mar;94(2):223–9.

27. Ginis H, Pérez GM, Bueno JM, Artal P. The wide-angle point spread function of the human eye reconstructed by a new optical method. J Vis. 2012 Mar;12(3):20, 1-10.

28. Cao D, Nicandro N, Barrionuevo PA. A five-primary photostimulator suitable for studying intrinsically photosensitive retinal ganglion cell functions in humans. J Vis. 2015 Jan;15(1):27, 1-13.

29. Barrionuevo PA, Cao D. Luminance and chromatic signals interact differently with melanopsin activation to control the pupil light response. J Vis. 2016 Sep;16(11):29, 1-17.

